# Triple COVID-19 vaccination induces humoral and cellular immunity to SARS-CoV-2 with cross-recognition of the Omicron variant and IgA secretion

**DOI:** 10.1101/2022.09.22.508999

**Authors:** Louisa Ruhl, Jenny F. Kühne, Kerstin Beushausen, Jana Keil, Stella Christoph, Jasper Sauer, Christine S. Falk

## Abstract

COVID-19 vaccination is the leading strategy to prevent severe courses after SARS-CoV-2 infection. In our study, we analyzed humoral and cellular immune responses in detail to three consecutive homologous or heterologous COVID-19 vaccinations. All individuals (n=20) responded to vaccination with increasing S1- /RBD-/S2-specific IgG levels, whereas specific plasma IgA displayed individual variability. The third dose increased antibody inhibitory capacity (AIC) against immune-escape variants Beta and Omicron independently from age. The mRNA-primed vaccination induced IgG and IgA immunity more efficiently, whereas vector-primed individuals displayed higher levels of memory T and B cells. Vaccinees showed a SARS-CoV-2-specific T cell responses, which were further improved and specified after Omicron breakthrough infections in parallel to appearance of new variant-specific antibodies. In conclusion, the third vaccination was essential to increase IgG levels, mandatory to boost AIC against immune-escape variants and induced SARS-CoV-2-specific T cells. Breakthrough infection with Omicron generates additional spike specificities covering all known variants.

## Introduction

The pandemic spread of the severe acute respiratory syndrome coronavirus 2 (SARS-CoV-2) has resulted in over 500 million infections with more than six million deaths due to the associated coronavirus disease 2019 (COVID-19). To contain the pandemic spread of SARS-CoV-2 and to prevent severe disease courses of COVID-19 an efficient immunity against the virus is crucial. To induce the latter, tremendous global efforts were taken to develop vaccines against SARS-CoV-2. Even though different vaccine-strategies were used, both mRNA- and vector-based vaccines were highly protective against viral infection [1, 2] and vastly effective in inducing humoral and cellular immune responses against SARS-CoV-2 after prime-boost vaccination [3, 4]. Notably, these vaccines were developed based on the spike-protein of the ancestral SARS-CoV-2 strain [5, 6]. Since then, several SARS-CoV-2 variants have emerged, challenging the immune response due to immune-escape mutations particularly in the spike-protein [7, 8]. Among those, the recent variant of concern (VOC) B.1.1.529 Omicron is evolutionary the most distant VOC to date [7, 9]. Omicron partially evades the humoral immune response, also in the mucosa in double vaccinated and convalescent individuals due to the spike mutations [10] and partly because protective antibody titers seem to decline over time [11, 12]. Therefore, booster (3^rd^) vaccinations are thought to induce recall immunity and to increase protection against VOC. The availability of different vaccine-platforms like mRNA- or vector-vaccines led to the application of different vaccine regimens. However, differences regarding the kinetics of immune responses between homologous *vs*. heterologous vaccinations need to be studied in more detail across adult age groups. In addition to specific antibody development, an effective T cell responses need be induced after COVID-19 vaccination as a second line of the adaptive defense. Particularly in the case of immune-escape variants like Omicron, the eventual lack of neutralizing antibodies requires a fast T cell response with primed SARS-CoV-2-specific T cells upon vaccination [13, 14]. Moreover, virus-specific T cells seem to be more durable and could compensate the waning humoral immune responses [15].

Our longitudinal matched study provides detailed insights into the fine tuning of specific immune responses after triple COVID-19 vaccination and their effectiveness against VOC in healthy, naïve individuals. We could demonstrate that a third vaccination is essential to increase spike-specific plasma IgG levels and, most importantly, to boost antibody inhibitory capacity (AIC) against Omicron and other VOC also in elderly vaccinees. Vaccination also induced spike-specific IgA secretion, indicating individual mucosal immunity against SARS-CoV-2. Moreover, spike-specific T cells were sufficiently induced after vaccination and developed into memory CD8^+^ and CD4^+^ T cells. Of note, the T cell response was lower in vaccinees compared to individuals with omicron breakthrough infections, who also displayed novel VOC-specific IgG. Taken together, our study demonstrates the benefits of three COVID-19 vaccinations for sustained immunity against SARS-CoV-2 and VOC and the capacity of the antibody repertoire to generate novel VOC-specific spike antibodies upon omicron infection even after triple vaccination with the wildtype sequence.

## Results

### Increasing levels of spike-specific IgG and IgA antibodies and high antibody-inhibitory-capacity with cross-recognition of VOC after three COVID-19 vaccinations

To assess the humoral immune response to COVID-19 vaccination, we quantified spike S1-, RBD- and S2-specific IgG, IgA and IgM antibodies in plasma samples from n=20 naive donors after first, second and third vaccination (**Fig.1A, Suppl. Tab. 1, 2**) via Luminex-based multiplex assays. Pre-pandemic matched blood samples obtained before vaccination severed as control (pre) with two subjects who displayed S1-specific, cross-reactive antibodies originating presumably from previous infections with common cold coronaviruses (**Suppl. Fig. 1**). 12 to 22 days after first vaccination S1-, RBD- and S2-specific IgG antibodies were detectable in plasma of vaccinated individuals (**Fig. 1B**) with substantial variability, i.e., high- and low-responders, but independent from age (data not shown). 20 to 49 days after the second dose, specific IgG levels massively increased also in low responders (**Fig. 1B**). To evaluate the maintenance of these spike-specific antibodies in their plasma, we additionally collected blood samples six months after the second vaccination (n=16). As expected, IgG levels declined variably but, of note, were still significantly higher as compared to the first and, of course, before vaccination (**Fig. 1B, Suppl. Fig. 1)**. Further waning of spike-specific IgG in the blood of vaccinees was prevented by a third dose of a COVID-19 vaccine, which resulted in increased IgG concentrations against S1-, RBD- and S2-antigens. Interestingly, spike IgG levels after the third dose were comparable to those after second vaccination (**Fig. 1B**). After second and third vaccinations, all analyzed individuals were seropositive for spike-specific antibodies, even six months after the second dose (**Fig. 1B, Suppl. Fig. 1**).

**Fig. 1.**
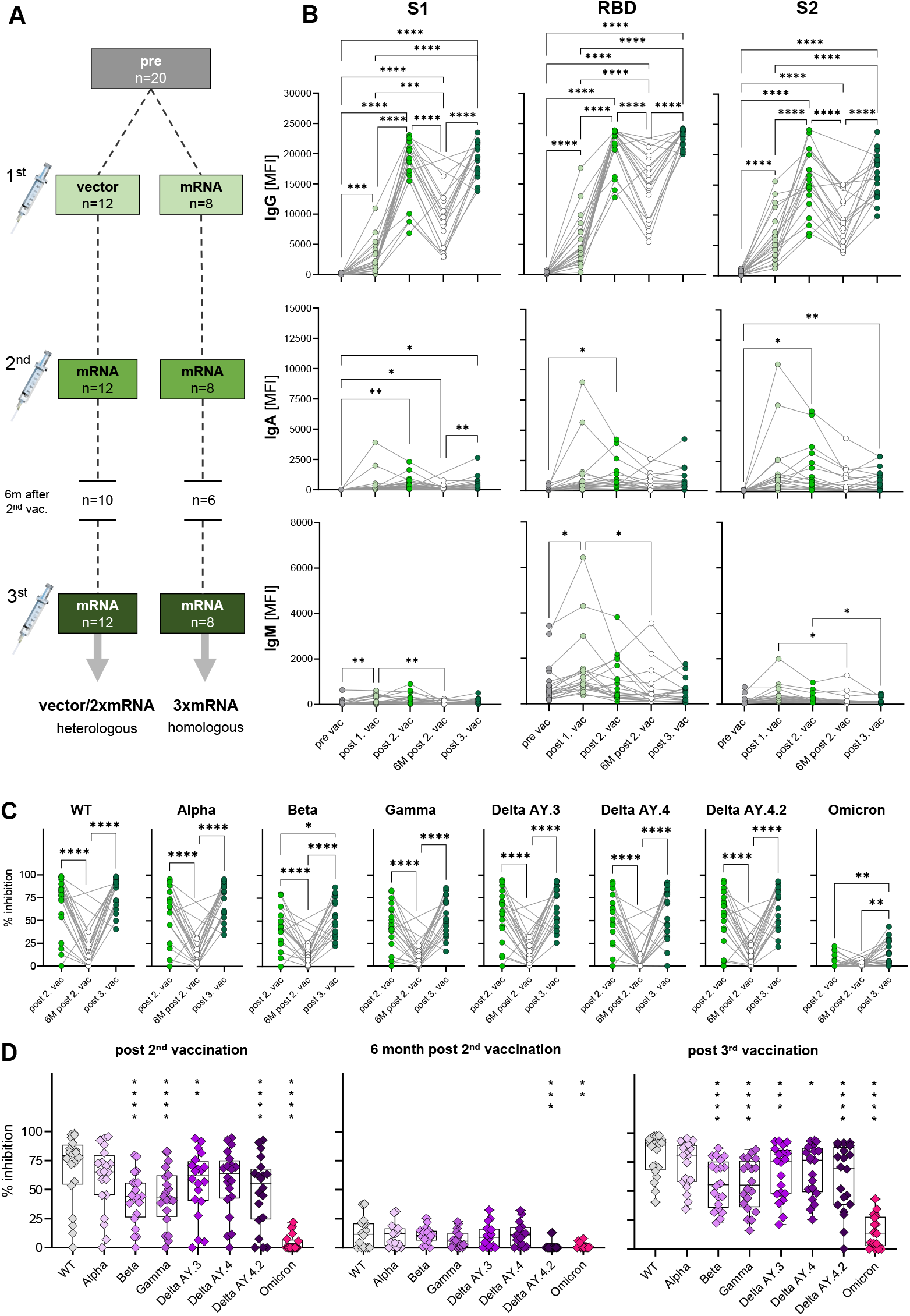
Antibody response and AIC against VOC to three consecutive vaccinations in healthy individuals. **(A)** Unexposed individuals were recruited to this study pre-vaccination (n=20), after first (n=20), second (n=20) and third (n=20) vaccination and six months after the second vaccination (n=16). Sample collection after vaccination is displayed as mean. **(B)** IgG, IgA and IgM antibody levels against SARS-CoV-2 S1-, S2-domain and RBD were measured with Luminex-based multiplex assays in n=20 individuals pre-vaccination, after first, second and third vaccination and in n=16 individuals six month after the second vaccination. Antibody levels are displayed as MFI. **(C, D)** Antibody inhibitory capacity (AIC) against several SARS-CoV-2 variants was analyzed using electrochemiluminescence-based multiplex assays and is displayed as % inhibition. **(C)** Comparison between AIC over time in vaccinees. **(D)** AIC against VOC (Alpha, Beta, Gamma, Delta AY.3, Delta AY.4, Delta AY4.4, Omicron) compared to WT after second vaccination, 6m after second vaccination and after third vaccination. Asterisks indicate p-value of significant differences between VOC and WT. Statistical analyses: (B, C) paired multi-group comparisons were performed using ANOVA test with Tukey multiple comparison test or (D) using Friedman test with Dunn’s multiple comparison test. *p < 0.05, **p < 0.01, ***p < 0.001, ****p < 0.0001. d: days, m: months, w: weeks.

In general, as IgA antibodies are predominantly present in mucosal tissues, their frequency in blood is naturally lower compared to IgG. IgA plasma levels against spike S1-, RBD- and S2-antigens varied highly between individuals but generally increased significantly after the second vaccination and after the third dose for S1- and S2-antigens (**Fig. 1B**). Nevertheless, we also observed highly increased IgA levels for some individuals already after first vaccination, arguing for a highly individual variance in class switch towards IgA.

Interestingly, in vaccines, we observed increased IgM levels especially after first vaccination (**Fig. 1B**). Particularly, RBD-specific IgM levels were higher compared to S1- and S2-specific IgM.

Concluding, COVID-19 vaccination was effective by inducing spike-specific humoral responses with highest IgG levels followed by IgA and IgM. Increasing levels of spike-specific IgA antibodies after vaccination indicated mucosal antibodies in at least some individuals conferring potential protection from infection.

The vaccines developed against COVID-19 in 2020 were based on the spike-protein sequence of the ancestral SARS-CoV-2 strain (WT). Since then, several VOC with different mutations in the spike-sequence emerged and spread globally. To investigate the antibody responses to COVID-19 vaccines not only quantitatively but also qualitatively, we performed antibody interference assays via electro-chemiluminescence-based multiplex assays, to determine whether antibodies of vaccinees were able to block *in vitro* binding of WT or VOC S1-domains to the human ACE2 receptor. This antibody-inhibitory capacity (AIC) is displayed as percent inhibition for each variant. In addition to WT, VOC were analyzed competitively in one well including B.1.1.7 (Alpha), B.1.351 (Beta), P.1 (Gamma), AY.3 (Delta), AY.4 (Delta), AY.4.2 (Delta) and B.1.1.529 (Omicron). To examine the AIC kinetics, matched samples after the second, six months after the second and after the third vaccination were analyzed. We observed high AIC against all VOC except Omicron already after the second vaccination that was significantly decreased after six months post second dose and restored with the third COVID-19 vaccine administration (**Fig.1C**). AIC after the second and third vaccination were similar for all VOC except for Beta and Omicron with significantly higher AIC after the third compared to second dose (**Fig. 1C**). These results underline the importance of a third COVID-19 vaccination for a neutralizing humoral immunity against different VOC. Moreover, IgG antibody levels were positively correlated with AIC against WT and all tested VOC after second and third vaccination (**Suppl. Fig. 2A, B**), indicating that high antibody titers result in high neutralizing capacity. The partial neutralization escape of certain VOC [7, 16] was confirmed by decreased AIC against the SARS-CoV-2 variants Beta, Gamma, Delta AY.3 and Delta AY.4 compared to WT (**Fig. 1D**). In summary, these results illustrate that the triple COVID-19 vaccination led to broad humoral immune responses that cross-recognized several SARS-COV-2 variants, but poorly Omicron.

### Homologous and heterologous vaccination groups after third vaccination do not differ in antibody levels and AIC

In our cohort, individuals were vaccinated with two different vaccine regimens and received either three doses of mRNA-vaccines (homologous, 3xmRNA, n=8) or one dose of an adenoviral-vector-vaccine (ChAdOx) followed by two doses of mRNA-vaccines (heterologous, vector/2xmRNA, n=12) (**Fig.1A, Suppl. Tab. 1, 2**). Both, homo- and heterologous vaccine regimens were compared regarding their IgG, IgA and IgM levels against S1-, RBD- and S2-spike domains. In general, the response to the two different vaccine regimens differed only after first and second vaccination (**Fig. 2A**). mRNA-primed individuals displayed higher levels of S1-, RBD- and S2-specific IgA antibodies after the first vaccination. In addition, S2-specific IgM were also increased in the homologous compared to the heterologous vaccinated cohort after first vaccination. After second vaccination, 3xmRNA-vaccinated individuals showed higher levels of RBD-specific IgG, whereas the vector/2xmRNA-vaccine combination led to increased levels of S2-specific IgG (**Fig. 2A**). Remarkably, these differences disappeared after the third vaccination, indicating that subtle priming effects of different vaccine regimens could be equalized by subsequent vaccinations. No differences in antibody responses were observed between heterologous and homologous vaccinations with regard to AIC against VOC-specific spike domains with lowest interference against Omicron (**Fig. 2B**). Taken together, slight differences were observed in the initial antibody levels between homologous and heterologous vaccine regimens, which were no longer visible after the third dose. Importantly, AIC against VOC was not affected by the vaccine regimens.

**Fig. 2.**
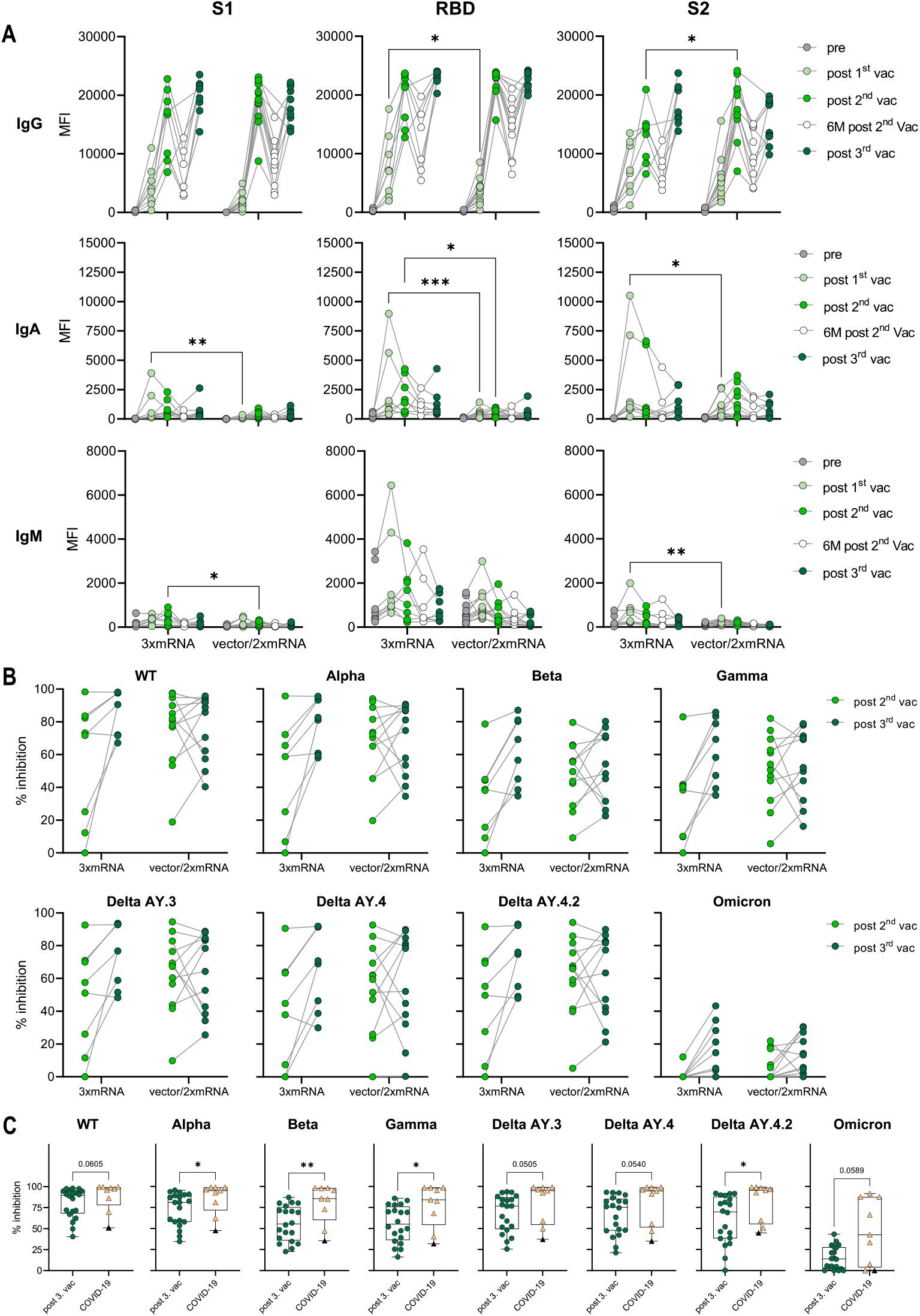
Antibody responses in homologous vs. heterologous and in vaccinated vs. infected individuals. **(A)** IgG, IgA and IgM antibody levels against SARS-CoV-2 S1-, S2-domains and RBD were measured with Luminex-based multiplex assays in n=20 individuals pre-vaccination, after first, second and third vaccination and in n=16 individuals six month after second vaccination. Antibody levels are displayed as MFI and were compared between homologous (3xmRNA) and heterologous (vector/2xmRNA) vaccine regimens. **(B, C)** Antibody inhibitory capacity (AIC) against several SARS-CoV-2 variants was analyzed using electrochemiluminescence-based multiplex assays and is displayed as % inhibition. **(B)** AIC against WT and VOC (Alpha, Beta, Gamma, Delta AY.3, Delta AY.4, Delta AY4.4, Omicron) after second and third vaccination was compared between homologous (3xmRNA, n=8) and heterologous (vector/2xmRNA, n=12) vaccine regimens. **(C)** AIC against WT and VOC (Alpha, Beta, Gamma, Delta AY.3, Delta AY.4, Delta AY4.4, Omicron) was compared between vaccinees after third dose (n=20) and infected individuals who were three-times vaccinated prior to breakthrough infection (COVID-19, n=9). Black triangle represents one individual who was infected with WT SARS-CoV-2 in 2021 and with Omicron in 2022 but unvaccinated. Statistical analyses: (A) paired multi-group comparisons were performed using ANOVA test with Tukey multiple comparison test. (B) Two-way repeated measures were performed using ANOVA test with Sidak multiple comparisons test. (C) Paired two-group comparison was performed using Mann–Whitney test; Spearman correlation. *p < 0.05, **p < 0.01, ***p < 0.001, ****p < 0.0001.

As the AIC against Beta and Omicron spike variants was strongly decreased compared to WT (**Fig. 1C, 1D**), humoral immunity after vaccination was also compared to breakthrough infections. Fully vaccinated and Omicron-infected individuals (COVID-19, n=8) with mild COVID-19 and one unvaccinated patient who was infected for twice, first with WT and second with Omicron were included. COVID-19 subjects showed higher AIC against the Alpha, Beta, Gamma and Delta VOC compared to vaccinees without infection (**Fig. 2C**). For the WT virus, we observed slightly increased AIC in the infected versus uninfected vaccinees. Moreover, infected individuals displayed higher AIC against Omicron compared to the uninfected vaccinated cohort. Nevertheless, the AIC against Omicron ranged from 0% to 91,6% in persons with breakthrough infection (**Fig. 2C**). Lowest AIC levels against all spike variants were obdserved in the non-vaccinated individual, suggesting that previous vaccination may improve antibody development upon SASR-CoV-2 infection. Interestingly, we observed that the vaccination cohort was split into two groups based on their AIC against all spike variants, arguing for high-vs. low-responders (**Fig. 2C**). As AIC strongly correlated with spike-specific IgG levels (**Suppl. Fig. 2A, B**), also high- and low-responders significantly differed in their IgG levels after third vaccination (**Suppl. Fig. 2C**). Notably, the age of low-responders did not diverge from high-responders (**Suppl. Fig. 2D**). Concluding, COVID-19 vaccination successfully induced antibodies capable to block the interaction between the SARS-CoV-2 S1-domain and the human ACE2 receptor. Nevertheless, vaccination-induced AIC was reduced for several VOC, especially Omicron compared to AIC after a breakthrough infection demonstrating the addition of new Omicron-specific antibodies.

### Differentiation of CD4^+^ T cells upon COVID-19 vaccination

To investigate the cellular immune response towards consecutive COVID-19 vaccinations, we performed an immunophenotyping of blood samples from n=19 unexposed donors after the first, second and third vaccination. Changes in the immunophenotype of vaccinated individuals were analyzed by quantifying absolute numbers of several leukocyte subsets via flow cytometry (**Suppl. Fig. 3**) and compared them with paired pre-vaccination samples. In general, we observed that immune cell numbers varied individually (**Fig. 3A-D**). After COVID-19 vaccination, B cells displayed dynamic changes in their phenotype, assessed by CD27 and IgD expression. Precisely, CD27^-^IgD^-^ double negative (DN) B cell numbers decreased over time after each vaccination. Simultaneously, CD27^+^IgD^+^ switch precursor B cell counts increased in response to first vaccination (**Fig. 3A**). Moreover, we observed a non-significant increase of CD27^+^IgD^-^ memory B cells and plasmablasts in some vaccinated individuals, especially after the first or third vaccination, respectively (**Fig. 3A**). Interestingly, B cell numbers slightly increased after first vaccination but declined subsequently after second and third dose for few individuals (**Fig. 3B**). No significant differences were observed between pre- and post-vaccination samples for granulocytes and lymphocytes, whereas monocyte counts were slightly decreased after second vaccination compared to pre-vaccination samples (**Fig. 3B**). T cell numbers were also rather stable and unaffected from vaccination, including CD4^+^ and CD8^+^ T cells (**Fig. 3B-D**). Expression of CCR7 and CD45RO was used to distinguish between naïve and memory cells. Numbers of CCR7^+^CD45RO^-^ naïve CD8^+^ T cells were marginally decreased after first and third vaccination compared to pre-vaccination samples (**Fig. 3C**). Simultaneously, numbers of CCR7^-^CD45RO^-^ TEMRA CD8^+^ T cells increased after first vaccination. Furthermore, a clear reduction of CCR7^+^CD45RO^+^ central memory (CM) CD8^+^ T cells in the periphery of vaccinated individuals was observed (**Fig. 3C**). Regarding CD4^+^ T cells, vaccinated donors displayed no differences in their number of naïve CD4^+^ T cells but, like for CD8^+^ T cells, numbers of CM CD4^+^ T cells were decreased after second and third vaccination compared to pre-vaccination controls (**Fig. 3D**). In addition, CCR7^-^CD45RO^+^ effector memory (EM) CD4^+^ T cells and TEMRA CD4^+^ T cells expanded after first vaccination (**Fig. 3D**). The latter also showed increased numbers after the third vaccination (**Fig. 3D**). Interestingly, we found numbers of CM CD4^+^ T cells to be negatively correlated to numbers of TEMRA CD4^+^ T cells after first and third vaccinations, arguing for a successful memory T cell development upon COVID-19 vaccination (**Suppl. Fig. 4**).

**Fig. 3.**
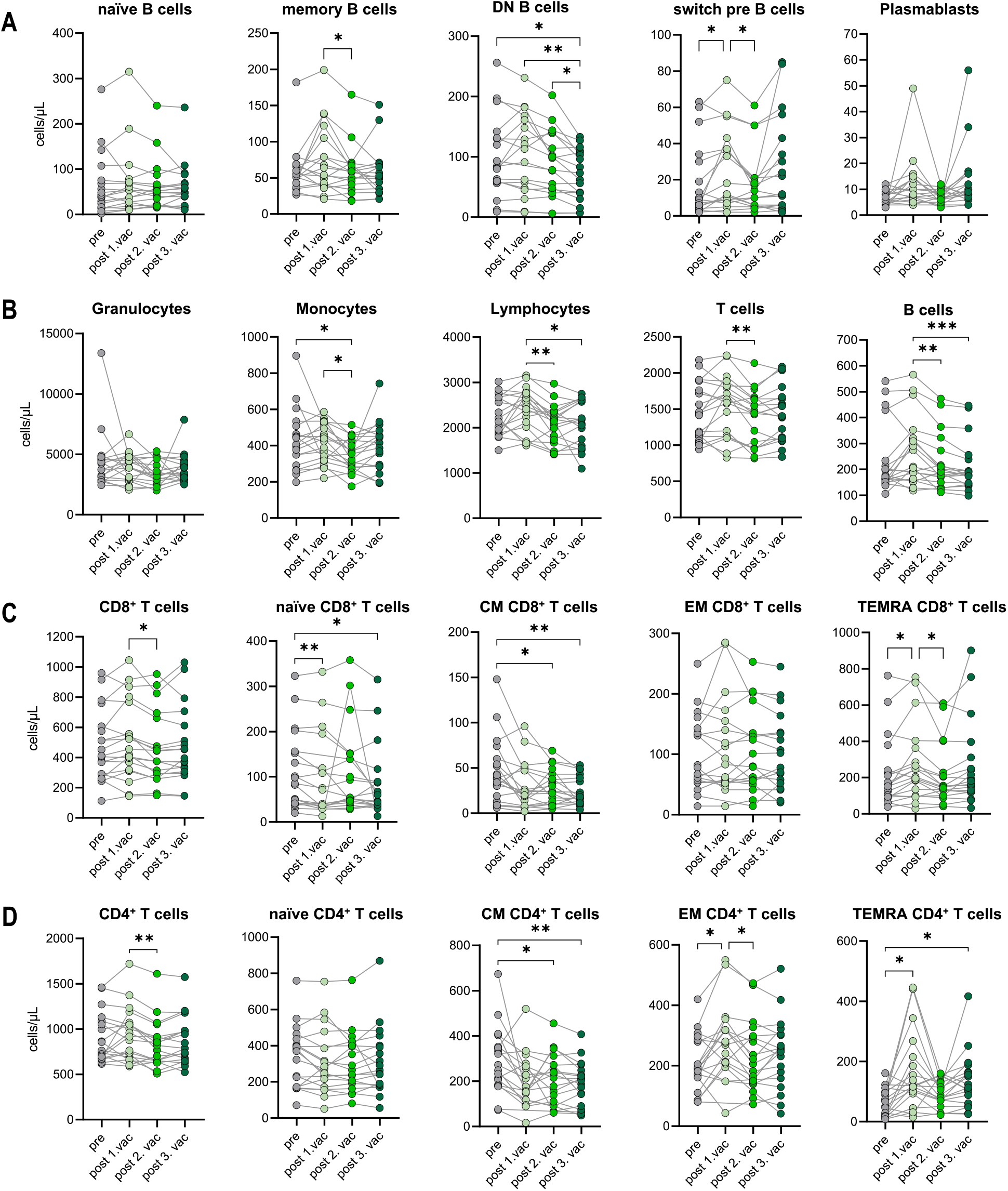
Immune cell phenotype before and after three consecutive vaccinations. Immune cell distribution presented by absolute numbers in blood was analyzed using Trucount analyses. Samples were analyzed pre-vaccination and after first, second and third vaccination in n=19 individuals. T cells: naive (CCR7^+^CD45RO^−^), central memory (CM, CCR7^+^ CD45RO^+^), effector memory (EM, CCR7−CD45RO^+^) and TEMRA (CCR7^−^CD45RO^−^); B cells: naive (IgD^+^CD27^−^) memory (mem, CD27^+^IgD^−^), switch precursor (switch pre, CD27^+^IgD^+^), double negative (DN, IgD^−^CD27^−^) and plasmablasts (CD19^+^CD20^−^CD27^+^CD38^+^). Gating strategy is shown in Supplementary Figure 3. CM: central memory, EM: effector memory, DN: double negative, switch pre: switch precursor. Statistical analyses: (A-D) paired multi-group comparisons were performed using ANOVA test with Tukey multiple comparison test. *p < 0.05, **p < 0.01, ***p < 0.001, ****p < 0.0001.

In summary, we found absolute numbers of T and B cell subsets to be highly individual and dynamic. Nevertheless, significant changes in the immunophenotype of unexposed donors after consecutive COVID-19 vaccinations were found. The reduction of CM with a simultaneous expansion of EM and TEMRA CD4^+^ and CD8^+^ T cells suggests a development of spike-specific memory T cells, which was supported by the negative correlation between CM and TEMRA CD4^+^ T cells counts.

### Subtle differences in T and B cell phenotype between mRNA- and vector-primed individuals

As we observed differences in the antibody response between homologous (3xmRNA) versus heterologous (vector/2xmRNA) vaccinated individuals (**Fig. 2A**), we determined the effect of different vaccine regimens (3xmRNA n=7; vector/2xmRNA n=12) on the immunophenotype regarding their T and B cell subsets. CD4^+^ and CD8^+^ T cell phenotypes were comparable between the 3xmRNA and the vector/2xmRNA vaccine groups. Only TEMRA CD4^+^ T cells were increased after first vaccination with a vector-vaccine, compared to a mRNA-vaccine (**Fig 4A, B**). A similar observation was made for memory B cells, which displayed higher numbers in heterologous compared to homologous vaccinated individuals after the first dose (**Fig. 4C**). For naïve, DN and switch precursor B cells, we found no differences between homologous and heterologous vaccine groups (**Fig. 4C**).

**Fig. 4.**
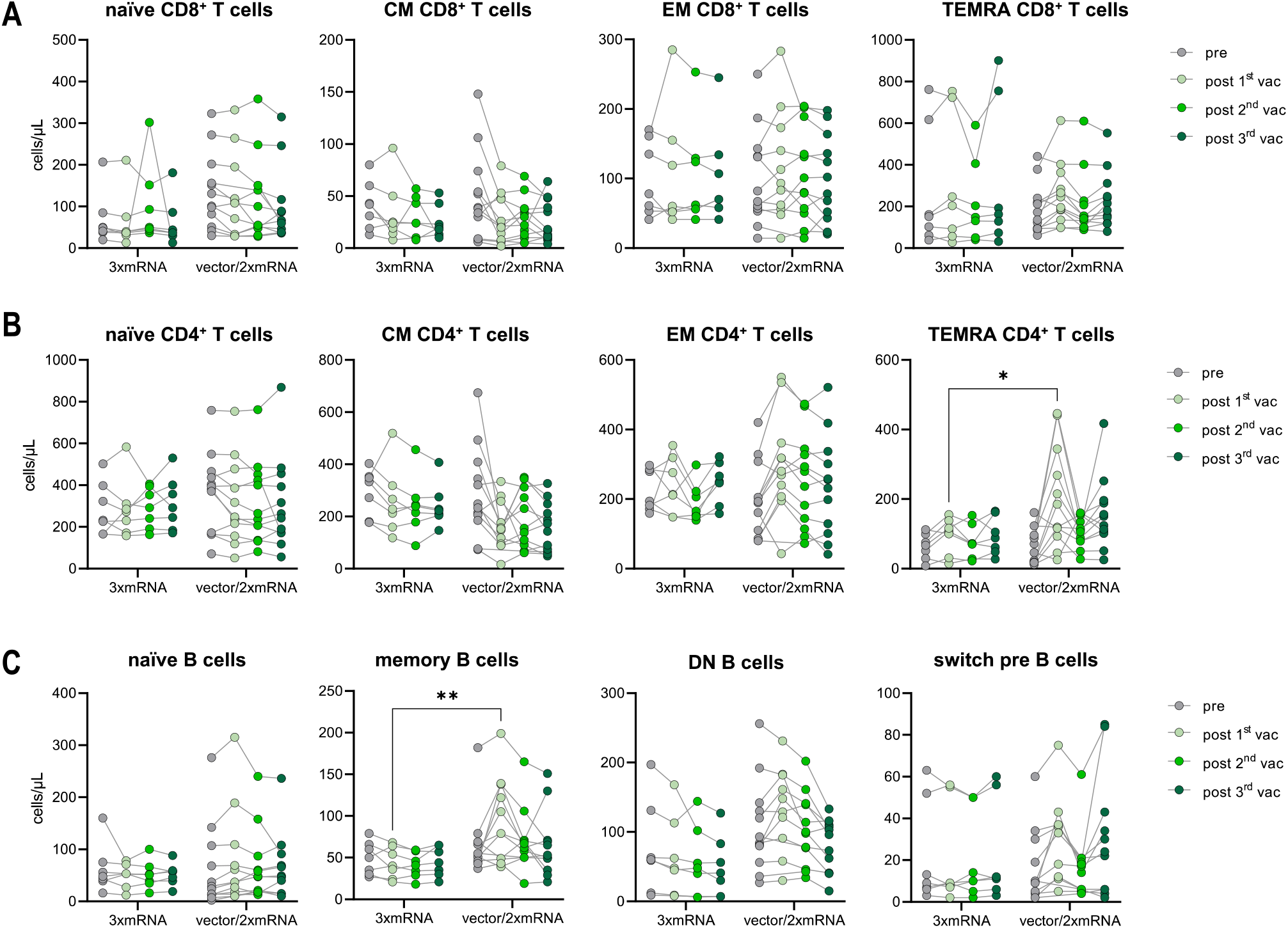
Immune cell phenotype in homologous vs. heterologous vaccinated individuals. Immune cell distribution presented by absolute numbers in blood was analyzed using TruCount analyses. Samples were analyzed pre-vaccination and after first, second and third vaccination and compared between homologous (3xmRNA, n=7) and heterologous (vector/2xmRNA, n=12) vaccine regimens individuals. T cells: naive (CCR7^+^CD45RO^−^), central memory (CM, CCR7^+^ CD45RO^+^), effector memory (EM, CCR7−CD45RO^+^) and TEMRA (CCR7^−^CD45RO^−^); B cells: naive (IgD^+^CD27^−^) memory (mem, CD27^+^IgD^−^), switch precursor (switch pre, CD27^+^IgD^+^), double negative (DN, IgD^−^CD27^−^) and plasmablasts (CD19^+^CD20^−^CD27^+^CD38^+^). Gating strategy is shown in Supplementary Figure 3. CM: central memory, EM: effector memory, DN: double negative, switch pre: switch precursor. Statistical analysis: (A-C) Two-way repeated measures were performed using ANOVA test with Sidak multiple comparisons test. *p < 0.05, **p < 0.01, ***p < 0.001, ****p < 0.0001.

These results imply an increased memory formation of CD4^+^ T cells and B cells after the first dose in individuals receiving an adenoviral-vector-vaccine (vector/2xmRNA) compared to mRNA-primed persons (3xmRNA). Importantly, differences in the immunophenotype were only found after first vaccination, indicating again that prime-effects of different vaccine regimens could be equalized by further vaccinations. Most importantly, there are no indications of sustained alterations in T and B cell subset compositions as consequence of vaccinations, which is in sharp contrast to COVID-19, especially severe disease courses [17].

### Effective but less broad T cell priming in COVID-19-vaccinated compared to SARS-CoV-2-infected individuals

To proof the development of spike-specific T cells *in vitro*, we investigated the T cell response by performing IFN-γ- and chemokine-release assays. Therefore, PBMC were stimulated with SARS-CoV-2 S1-antigens and the following T cell response was assessed by measuring several cytokines and chemokines associated with T cell activation in culture supernatant. T cell responses of vaccinated individuals were compared to those of triple vaccinated individuals with breakthrough infections (COVID-19, n=8). Paired, unstimulated samples served as control. Upon stimulation with S1-antigen, increased IFN-γ secretion along with Th1-associated cytokines and chemokines was observed in both cohorts, vaccination and breakthrough infection (**Fig. 5A, B**), suggesting effective memory T cell formation in both conditions. However, higher IFN-γ secretion was observed in individuals with breakthrough infections compared to vaccinated individuals without virus contact (**Fig. 5C**). Interestingly, no differences in T cell response were observed between homologous and heterologous vaccine cohorts (**Fig. 5D**). Despite IFN-γ, we analyzed several other cytokines and chemokines associated with T cell activation such as IL-1RA, IL-2, CXCL8, TNF-α, G-CSF, CCL3 and CCL4. While infected individuals also responded to S1-stimulation with increased secretion of IL-1RA, IL-2, CXCL8, TNF-α, G-CSF, CCL3 and CCL4, only a subgroup of vaccinated individuals did so (**Fig. 5A-C**). These results indicate a broader T cell response and probably improved T cell priming upon breakthrough infection compared to triple vaccination alone. Furthermore, our finding suggest that the T cell response is, in contrast to the humoral immune response, not affected by vaccine regimens.

**Fig. 5.**
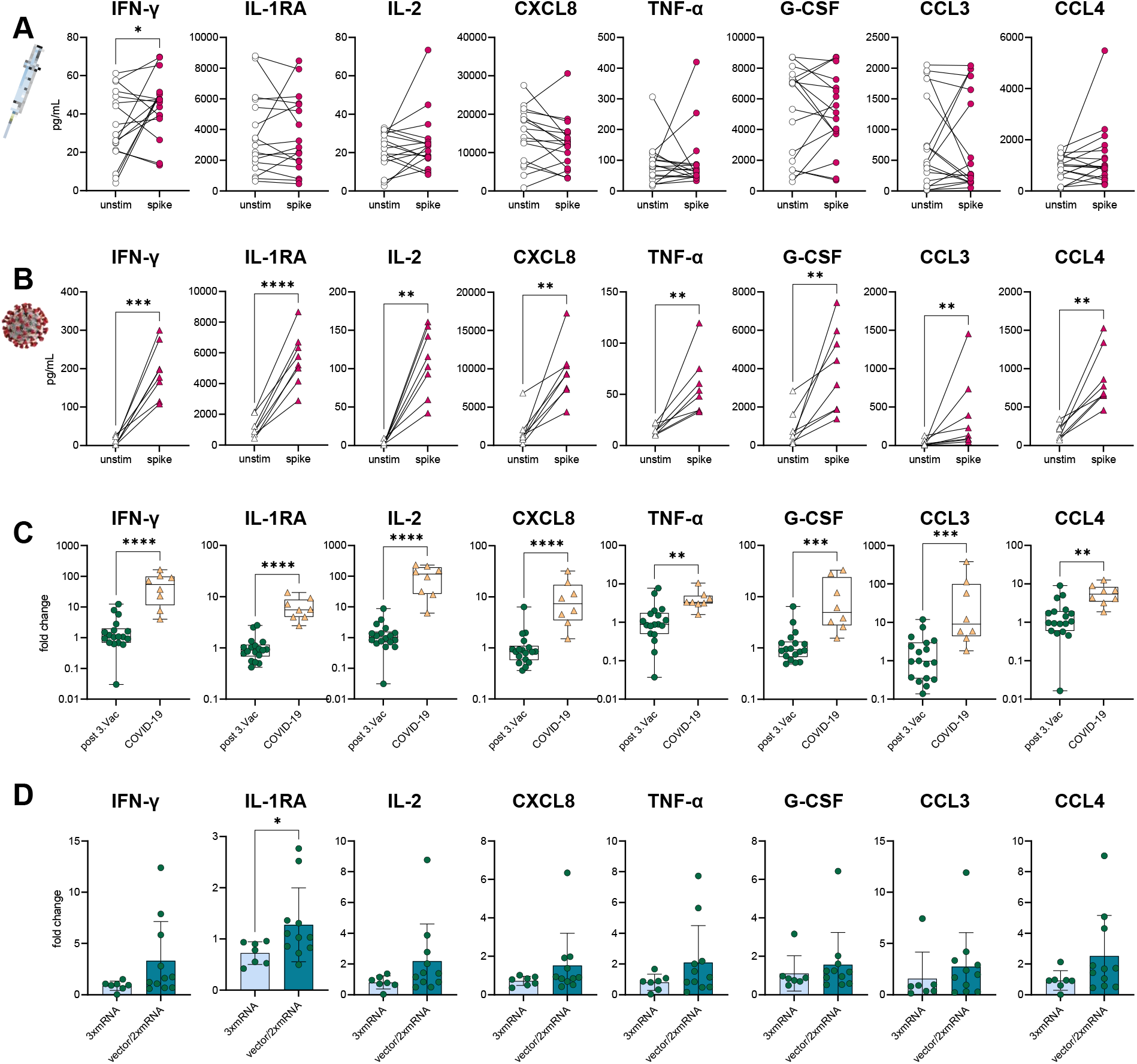
S1-specific T cells in vaccinees and infected individuals. SARS-CoV-2-specific T cell response was assessed in **(A)** vaccinated (n=18) and **(B)** infected individuals (COVID-19, n=8) by using IFN-γ and chemokine release assays based on *in vitro* stimulation of T cells with SARS-CoV-2 S1-antigens. Cytokines and chemokines were detected in culture supernatants (vaccinees) or plasma (COVID-19) using Luminex-based multiplex assays. Samples from vaccinated individuals were obtained post third vaccination. **(A, B)** T cell response was assessed by comparing cytokine and chemokine secretion from unstimulated versus S1-stimulated samples. **(C)** Comparison between vaccinated and infected individuals regarding their S1-specific T cell response. T cell response was displayed as fold change (stimulated/unstimulated). **(D)** Comparison of the S1-specific T cell response between homologous (3xmRNA, n=6) and heterologous vaccine regimens (vector/2xmRNA, n=11). T cell response was displayed as fold change (stimulated/unstimulated). Statistical analyses: (A, B) Paired two-group comparison was performed using t-test or Wilcoxon test. (C, D) Two-group comparison was performed using t-test or Mann-Whitney test. *p < 0.05, **p < 0.01, ***p < 0.001, ****p < 0.0001.

## Discussion

The COVID-19 vaccination is the leading strategy to overcome the worldwide pandemic with more than 500 million infections and over six million deaths caused by SARS-CoV-2 and should prevent severe disease progression after infection. Preclinical prime boost studies of mRNA-[1] and vector-based vaccine candidates [2] proofed efficient induction of spike-specific humoral and cellular immune responses. However, since the successful development of COVID-19 vaccines, several VOC with a variety of mutations emerged. Especially the appearance of immune-escape-variants, like Beta and Omicron, challenge the vaccine-induced immune response and led to a recommended vaccine regimen of three COVID-19 vaccinations. With the currently spreading immune-escape-variants and the high likelihood of further VOCs evolving in the future, it becomes even more important to study and to understand the vaccine-induced immunity to SARS-CoV-2 after at least three COVID-19 vaccine doses. Therefore, we aimed to broadly investigate the SARS-CoV-2-specific immune response including immunophenotyping, humoral and T cell responses to three consecutive COVID-19 vaccinations in n=20 healthy individuals.

Our data demonstrate that the levels of IgG, IgA and IgM antibodies against three different SARS-CoV-2 spike-antigens dynamically changed over time after each vaccination, which was also reported by other studies. The increase in IgG levels after second dose proofed the importance of an additional antigen exposure for the humoral immune response [12, 18, 19]. The reduction of antibody levels over time [11, 12] was impeded by a third vaccination, leading to a strong increase in plasma IgG levels. Moreover, our data revealed the importance of the third vaccination for antibody development against the immune-escape variants Beta and Omicron, since AIC against these VOC was significantly elevated after a booster dose compared to second vaccination. However, even after third vaccination the AIC against Omicron did not increase above 50% in vaccinated individuals, suggesting an at least partial antibody-evasion of the Omircon variant. The immune-escape by Omicron has been shown to be caused by numerous mutations, especially in the RBD, leading to decreased antibody neutralization potency [7]. Nevertheless, Omicron-neutralizing memory B cells were found in vaccinated individuals and provided at least some protection against immune-escape variants, even though their portion was shown to be reduced [20]. We observed a difference in AIC against Beta and Omicron after second and third vaccination, which could be explained by the reported development of new antibody clones upon booster vaccination that particularly target more conserved region of the RBD [12]. Consequently, these antibodies are more efficient in neutralizing highly mutated SARS-CoV-2 variants as Beta and Omicron. This proofs a strong but incomplete immune evasion by the Omicron variant and leads to the conclusion, that the current vaccines based on the ancestral strain supposedly are capable to elicit a variant-specific immune response.

In our study, breakthrough infection with Omicron led to increased AIC against this variant and other VOC compared to three-times vaccinated individuals, which could be explained by a recall of memory B cells that cross-recognized Omicron due to shared epitopes among SARS-CoV-2 variants [16]. Interestingly, a breakthrough infection with Omicron BA.1 was shown to induce a strong neutralization activity against BA.1 and BA.2 but not against Omicron subtypes BA.4 and BA.5 [16]. Additionally, triple mRNA-vaccinated individuals were characterized by a decreased capacity to neutralize Omicron BA.4 and BA.5 compared to BA.1 [16], suggesting a further immune-escape of the newly emerging Omicron subtypes.

As the SARS-COV-2 entry- and infection-pathway primarily involves the respiratory tract with mucosal tissue, the mucosal immunity mediated by tissue-resident T cells and IgA antibodies becomes of particular interest. Notably, expansion of nasal tissue-resident CD69^+^CD103^+^CD8^+^ T cells were detected after mRNA vaccination [21] and, as others and we have shown [12], SARS-CoV-2-specific IgA antibodies increased in plasma after vaccination. These data indicate a development of tissue-localized humoral and cellular mucosal immunity after intramuscular COVID-19 vaccination.

To fight the SARS-CoV-2 pandemic, several COVID-19 vaccines based on different vaccine platforms were developed, e.g., adenoviral-vector-based or mRNA-based vaccines. The availability of different COVID-19 vaccines led to numerous various vaccine combinations, raising the question about the immune response to homologous *vs*. heterologous vaccine regimens. We observed that heterologous and homologous vaccinated individuals displayed slight differences in their humoral immune response after first and second vaccination. Similar to other reports, we observed that the mRNA-primed vaccine regimen induced higher levels of RBD-specific IgG and S1-, RBD- and S2-specific IgA antibodies [22-26]. On the other hand, the vector-primed vaccination seemed to induce a more potent cellular response as seen by elevated numbers of TEMRA CD4^+^ T and memory B cells. In line with this, Schmidt et al. found spike-specific CD69^+^IFN-γ^+^ CD4^+^ T cells to be increased in vector-vaccinated compared to mRNA-vaccinated individuals [17]. Interestingly, after second vaccination with an mRNA-based vaccine, frequencies of spike-specific CD69^+^IFN-γ^+^ CD4^+^ T cells were comparable between vector- and mRNA-primed individuals, which is consisted with our observations where we found no differences in T cell numbers and cytokine secretion between homologous and heterologous vaccine regimens after second vaccination. Independent from the vaccine regimens, we observed a continued decline in peripheral CM CD4^+^ and CD8^+^ T cell numbers after each vaccination, arguing for a sustain differentiation of memory T cells. Because of waning antibody levels after vaccination and antibody neutralization resistance of immune-escape variants, an efficient T cell mediated immune response is crucial. In our study, we observed spike-specific IFN-γ producing T cells in vaccinated individuals, proving the development of spike-specific T cells after vaccination. Nevertheless, individuals with a breakthrough infection displayed a broader T cell response with secretion of multiple cytokines and chemokines upon spike re-stimulation compared to vaccinees. Importantly, the infected individuals were three-times vaccinated prior to their breakthrough infection. This multiple antigen exposure could explain the broader T cell response in vaccine-primed infected individuals. In line with this, Lang-Meli et al. demonstrated that in naïve individuals the T cell response was similar between second and third vaccination, whereas convalescent individuals benefited from a post-infection vaccination with an elevated SARS-CoV-2-specific T cell response [27]. However, in contrast to the humoral immune response, SARS-CoV-2-specific T cells seem to be more durable and efficiently cross-recognize the Omicron variant in vaccinated individuals [14, 15, 28]. These observations underline the importance of T cells for an effective virus-specific immune response and protective immunity against emerging VOC in vaccinees.

In conclusion, we observed that the triple COVID-19 vaccination is highly effective and induces humoral as well as cellular immune responses. Besides IgM and IgA, high concentrations of spike-specific IgG antibodies were produced after vaccination. Therefore, the level of IgG antibodies strongly correlated with AIC against several VOC and could distinguish high- and low-responder. For inhibition of immune-escape variants as Beta and Omicron the third vaccination seems to be crucial, since it boosted the AIC compared to second vaccination. The immunophenotype of vaccinees revealed a positive vaccination effect on cellular level with expansion of memory T and B cells and the development of S1-specific IFN-γ secreting T cells. The homologous and heterologous vaccine regimens displayed no differences regarding the humoral or cellular immune response after third vaccination. However, for the first vaccination mRNA-priming induced a stronger humoral and vector-priming an increased cellular immune response.

## Limitationsof the study

The limitations of our observational study include the single-center setting with a rather small sample size and time-points for sample collection were not standardized.

## Material & Methods

### Study design

In total 20 vaccinated individuals and 9 persons with breakthrough infections were recruited to this observational study between May 2020 and February 2022 at the Hannover Medical School (MHH, ethical vote 9001_Bo_K). Samples from vaccinated individual were collected before vaccination, after first and second vaccination, six months after second vaccination and after third vaccination. Demographical characteristics of study participants are summarized in Supplementary Table 1. Vaccinated individuals received either a homologous vaccination consisting of three-times mRNA vaccines (3xmRNA) or a heterologous vaccination with an adenoviral-vector-vaccine followed by two doses of mRNA vaccines (vector/2xmRNA, Supplemental Tab. 2). Infected subjects had a breakthrough infection with the Omicron variant and were three-times vaccinated at the time of infection, expect for one person who was not vaccinated but previously infected with SARS-CoV-2 WT.

### Multiplex assays

Luminex-based multiplex assays were used to quantify cytokines and chemokines as well as SARS-CoV-2 S1-, RBD- and S2-specific antibodies. Cytokines and chemokines were measured using the Bio-Plex Pro™ Human Assay (Bio-Rad, Hercules, USA): cytokine screening panel plus ICAM-1 and VCAM-1 (12007283, 171B6009M, 171B6022M) following manufacturer’s instructions. As samples thawed supernatant or plasma, which were diluted twofold with assay buffer were used. Standards were reconstituted and prepared as described in the manufacturer’s instructions. Standard curves and concentrations were calculated using the Bio-Plex Manager 6.1 software.

SARS-CoV-2 S1-, RBD- and S2-specific antibodies were detected using the SARS-CoV-2 Antigen Panel 1 IgG, IgM, IgA assay (Millipore, HC19SERM1-85K-04, HC19SERA1-85K-04, HC19SERG1-85K-04) following manufacturer’s instructions. Thawed plasma were used and diluted 1:100 with assay buffer.The semi-quantitative readout is given as median fluorescence intensity (MFI) of > 50 beads for each antigen and sample, acquired by the Bio-Plex 200 machine and the Bio-Plex Manager™ Version 6.0 software (Bio-Rad, Hercules, USA). BAU standard curve was generated by measuring calibrators of the Anti-SARS-CoV-2-QuantiVac-ELISA (Euroimmun, Germany, EI 2606-9601-10 G). BAU values were calculated using the Bio-Plex Manager 6.1 software based on the BAU standard curve.

### Electrochemiluminescence multiplex assays

For analyzing SARS-CoV-2 variant-specific antibody inhibitory capacity (AIC) a multiplex serology assay (V-PLEX SARS-CoV-2 Panel 23 (IgG), K15567U, Mesoscale, USA) for IgG antibodies to eight spike antigens from variants of SARS-CoV-2, including the (Alpha), B.1.351 (Beta), P.1 (Gamma), AY.3 (Delta), AY.4 (Delta), AY.4.2 (Delta) and B.1.1.529 (Omicron) variants was used. As samples thawed plasma which was diluted 1:100 with assay buffer was used. The assay was performed according to manufacturer’s instructions and samples were acquired by the MESO QuickPlex SQ 120. AIC was calculated using the MSD Discovery Workbench Software and visualized as % inhibition.

### Quantification of cells from EDTA blood via Trucount™ analysis

BD Trucount™ Tubes (BD Biosciences) were used to calculate absolute cell numbers from whole blood following manufacturer’s instructions.

### Flow cytometry

Flow cytometry analyses were performed as recommended by the guidelines of leading European scientists of immunology and flow cytometry communities [29]. 100 µL whole blood EDTA samples were incubated with antibodies for surface staining in FACS Buffer (0,1% NaN3, 2,5% FCS in PBS) at 4°C for 30 min and followed by 15 min erythrocyte lysis using 1x BD Lysing Solution. Prior to acquisition cells were washed with PBS. All antibodies used for flow cytometry analyses are listed in Supplementary Table 3. Cells were acquired and analyzed on a LSRII flow cytometer (BD Biosciences, USA) using FACS Diva software (v8.0).

### IFN-γ and chemokine release assay

The IFN-γ and chemokine release assay “Quan-T-Cell SARS-CoV-2” (Euroimmun, Germany) was used to detect spike S1-specific T cells. For vaccinated individuals we performed the assay the PBMC and for infected individuals we used heparin whole blood samples. PBMC were thawed and seeded with 7×10^5^ to 1×10^6^ per tube in 500 µl medium (RPMI1640 +2mM L-Glutamin + 100 U/mL Penicillin-Streptomycin +1Mm sodium pyruvate +10% FCS) or 500 µl heparin-whole blood was added to each tube. Subsequently, tubes were inverted and incubated for 20 h at 37°C with 5% CO2. After the incubation, supernatant or plasma was collected by centrifuging the tube for 10 min at 6000g. The supernatant and plasma were frozen until further usage. Cytokines and chemokines in the supernatant or plasma were measured via Luminex-based multiplex assays.

### Statistical analyses

Statistical analyses of the data were performed with GraphPad Prism v9.0 software (GraphPad Software). To assess data distribution, Anderson-Darling normality test was calculated. Parametric tests were performed where data were normally distributed, otherwise non-parametric tests were used. The statistical tests used in each analysis are indicated in the figure legends. Correlation analyses were performed using Spearman-rank-order correlation. Results were considered significant if p<0.05.

### Study approval

The study was approved by the Hannover Medical School Ethics Committee. All patients or participants provided written informed consent before participation in the study (9001 BO K, 968-2011).

### Data availability

The datasets used and/or analyzed to support the findings of this study are available in this paper or the Supplementary Information. Any other raw data that support the findings of this study are available from the corresponding author upon reasonable request.

## Acknowledgments

This project was supported by the German Research Foundation DFG FA-483/1-1, the German Center for Infection Research DZIF TTU-IICH 07_913, the Lower Saxony Ministry of Research and Culture (ImProVIT) and the COFONI fast track project 8FT21.

## Conflict of interest statement

The authors have declared that no conflict of interest exists.

## Author contribution

LR performed flow cytometry experiments as well as Luminex-based and electrochemiluminescence-based multiplex assays, analyzed data, prepared figures and wrote the manuscript. JFK performed flow cytometry experiments and supported writing the manuscript. KB and JK performed Luminex-based and electrochemiluminescence-based multiplex assays. SC and JS collected blood samples. CSF supervised and designed the study and wrote the manuscript. All authors have read and approved the article.

**Supplementary Fig. 1.**
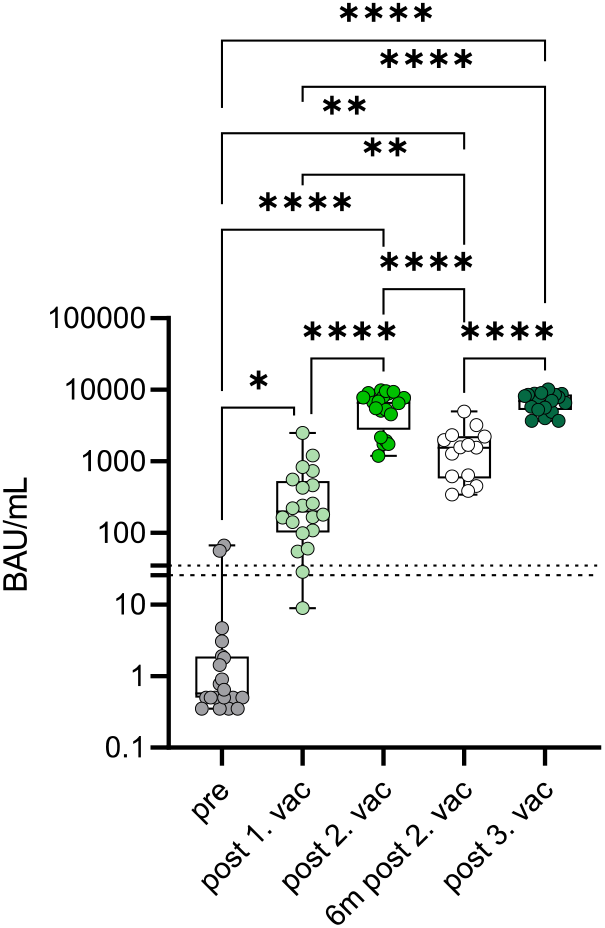
Antibody levels displayed as BAU/mL. Luminex-based multiplex assays were used to quantify SARS-CoV-2 S1-specific IgG antibodies in n=20 individuals pre-vaccination, after first, second and third vaccination and in n=16 individuals six month after second vaccination. Calibrators of the Anti-SARS-CoV-2-QuantiVac-ELISA were used to generate the BAU standard curve. BAU values were calculated using the Bio-Plex Manager 6.1 software. Cut off of >35,2 BAU/mL was defined as seroconverted and is represented by the dotted line. BAU/mL values between 25,6-35,2 are considered to be marginal, displayed by the dashed line. Statistical analyses: multi-group comparisons were performed using ANOVA test with Tukey multiple comparison. *p < 0.05, **p < 0.01, ***p < 0.001, ****p < 0.0001.

**Supplementary Fig. 2.**
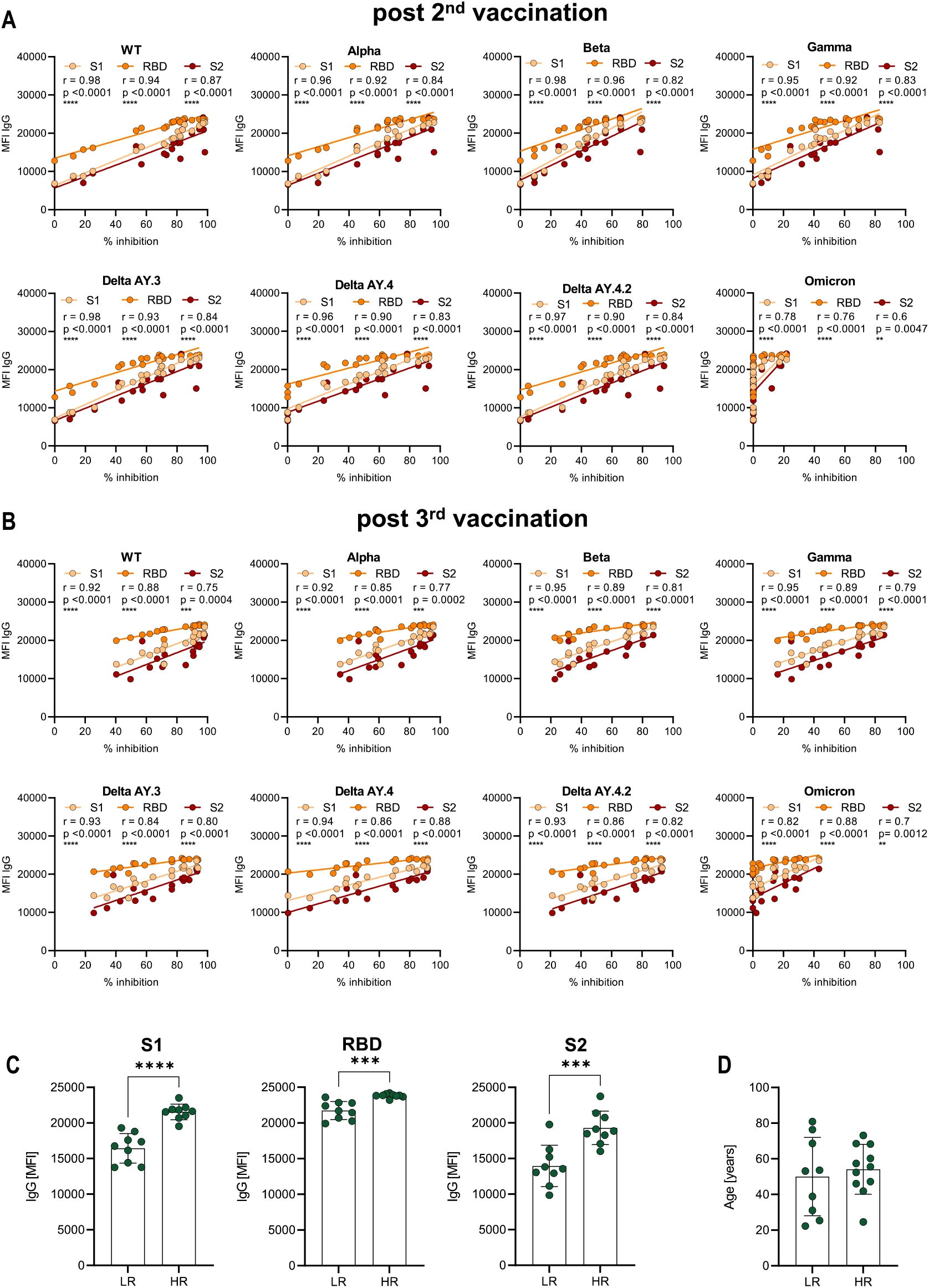
Correlation between IgG antibody levels and AIC against different SARS-CoV-2 variants. Luminex-based multiplex assays were used to quantify SARS-CoV-2 S1-, S2- and RBD-specific IgG antibodies in n=20 individuals after second **(A)** and third **(B, C)** vaccination. Antibody inhibitory capacity (AIC) against several SARS-CoV-2 variants was analyzed using electrochemiluminescence-based multiplex assays and is displayed as % inhibition. AIC was correlated to IgG levels after second **(A)** and third **(B)** vaccination. **(C)** Comparison of S1-, S2- and RBD-specific IgG antibody levels between low- (LR) and high-responders (HR). High responders were defined to have an AIC ≧mean for at least five of the analyzed VOC. **(D)** Age comparison between LR and HR. Statistical analyses: (A, B) Correlation analyses was performed using Spearman Rank correlation. (C, D) Two-groups comparison was performed using unpaired t-test. *p < 0.05, **p < 0.01, ***p < 0.001, ****p < 0.0001.

**Supplementary Fig. 3.**
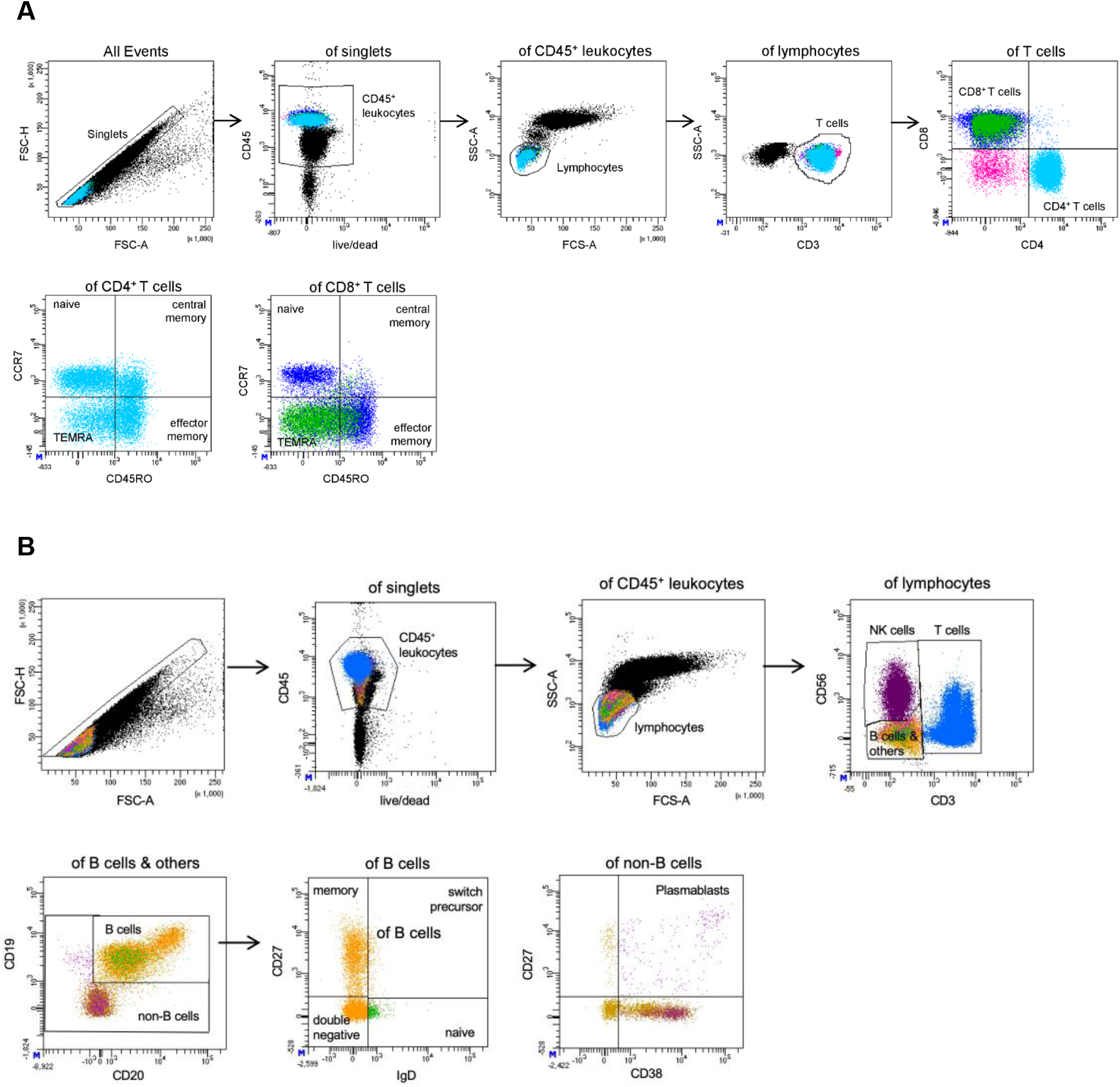
Flow cytometry gating strategy. Representative flow cytometry plots visualizing the gating strategy for **(A)** T cell subsets and **(B)** B cell subsets.

**Supplementary Fig. 5.**
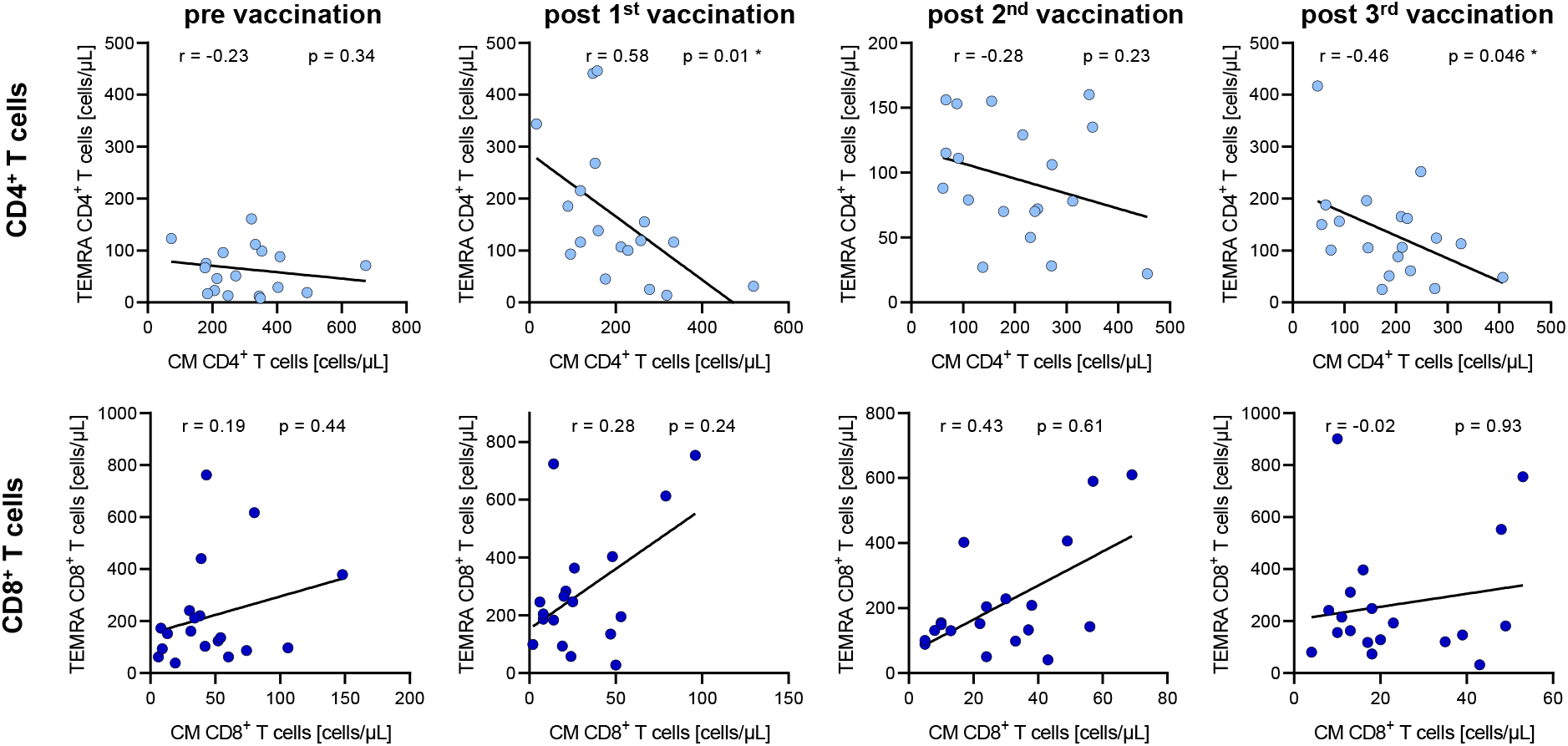
Correlation analyses between CM and TEMRA CD4^+^ and CD8^+^ T cells. CD4^+^ **(A)** and CD8^+^ T cell **(B)** distribution presented by absolute numbers in blood was analyzed using Trucount analyses. Samples were analyzed pre-vaccination and after first, second and third vaccination in n=19 individuals. Gating strategy is shown in Supplementary Figure 3. **(A)** Correlation analysis between CM and TEMRA CD4^+^ T cell numbers pre-vaccination and after first, second and third vaccination. **(B)** Correlation analysis between CM and TEMRA CD8^+^ T cell numbers pre-vaccination and after first, second and third vaccination. CM: central memory (CCR7^+^CD45RO^+^), TEMRA (CCR7^−^CD45RO^-^). Statistical analyses: Correlation analysis were performed using Spearman Rank correlation. *p < 0.05, **p < 0.01, ***p < 0.001, ****p < 0.0001.

**Table S1:** Demographics of vaccinated individuals and infected persons. Characteristics of individuals who were three-times vaccinated or SARS-CoV-2 infected people (COVID-19). Vaccinated individuals received either with a homologous vaccination of a mRNA vaccine (3xmRNA) or a heterologous vaccination with an adenoviral vaccine plus two-time mRNA (vector/2xmRNA). n=number of donors.

| Characteristics | 3xmRNA cohort | vector/2xmRNA cohort | COVID-19 cohort |
| --- | --- | --- | --- |
| Sample number n | 8 | 12 | 9 |
| Age in years (Min - Max) | 68,8<br>(46 - 80,9) | 42,2<br>(22,3 - 60,2) | 26,5<br>(23,6 - 29,3) |
| Gender |  |  |  |
| Female | 3 (43%) | 9 (75%) | 6 (66,7%) |
| Male | 4 (57%) | 3 (25%) | 3 (33,3%) |
| Time past after Vac. |  |  |  |
| Days post 1st Vac. | 20 | 14 | n.A |
| Days post 2nd Vac. | 29 | 23 | n.A |
| Weeks post 3rd Vac. | 7 | 5 | n.A |
| Time past after infection |  |  |  |
| Weeks post infection | n.A | n.A | 4,8 |

**Table S2:** Demographics and vaccine information of each vaccinated individual. Individuals demographics of triple vaccinated individuals. Vaccinated individuals received either with a homologous vaccination of a mRNA vaccine (3xmRNA) or a heterologous vaccination with an adenoviral vaccine plus two-time mRNA (vector/2xmRNA). ChAd: ChAdOx-1 vaccine, BNT: BNT162b2 vaccine, MDN: mRNA-1713 vaccine, d: days, w: weeks

| Nr. | Age | Sex | 1st vaccine | 2nd vaccine | 3rd vaccine | time post 1st vac | time post 2nd vac | time post 3rd vac | vaccine cohort |
| --- | --- | --- | --- | --- | --- | --- | --- | --- | --- |
| 1 | 53,6 | F | ChAd | BNT | BNT | 14d | 20d | 4W | vector/2xmRNA |
| 2 | 52,4 | M | ChAd | BNT | BNT | 13d | 21d | 6W | vector/2xmRNA |
| 3 | 80,9 | F | BNT | BNT | BNT | 3W | 7W | 7W | 3xmRNA |
| 4 | 60,2 | M | ChAd | MDN | MDN | 18d | 23d | 6W | vector/2xmRNA |
| 5 | 24,5 | F | ChAd | BNT | BNT | 14d | 27d | 5W | vector/2xmRNA |
| 6 | 68,5 | M | BNT | BNT | BNT | 17d | 25d | 7W | 3xmRNA |
| 7 | 31,0 | M | ChAd | BNT | BNT | 12d | 26d | 8W | vector/2xmRNA |
| 8 | 38,9 | F | ChAd | BNT | MDN | 15d | 27d | 5W | vector/2xmRNA |
| 9 | 55,5 | F | ChAd | BNT | MDN | 14d | 21d | 5W | vector/2xmRNA |
| 10 | 53,1 | F | ChAd | BNT | MDN | 13d | 20d | 5W | vector/2xmRNA |
| 11 | 68,3 | M | BNT | BNT | BNT | 16d | 24d | 7W | 3xmRNA |
| 12 | 25,4 | F | ChAd | BNT | BNT | 13d | 22d | 4W | vector/2xmRNA |
| 13 | 73,1 | F | BNT | BNT | MDN | 18d | 28d | 5W | 3xmRNA |
| 14 | 68,6 | F | BNT | BNT | BNT | 14d | 21d | 7W | 3xmRNA |
| 15 | 57,4 | M | BNT | BNT | BNT | 14d | 3W | 4W | 3xmRNA |
| 16 | 46,0 | M | BNT | BNT | MDN | 29d | 24d | 7W | 3xmRNA |
| 17 | 76,4 | M | BNT | BNT | BNT | 22d | 32d | 8W | 3xmRNA |
| 18 | 22,3 | F | ChAd | BNT | BNT | 13d | 23d | 8W | vector/2xmRNA |
| 19 | 47,2 | F | ChAd | BNT | BNT | 16d | 29d | 5W | vector/2xmRNA |
| 20 | 41,9 | F | ChAd | BNT | MDN | 13d | 21d | 4W | vector/2xmRNA |

**Table S3:** Flow cytometry antibodies. Antibodies used for flow cytometry analyses.

| Antigen | Fluorochrome | Manufacturer |
| --- | --- | --- |
| CD3 | V500 | BD Bioscience |
| CD3 | APC-H7 | BD Bioscience |
| CD3 | PerCP | BD Bioscience |
| CD4 | PerCP | BD Bioscience |
| CD6 | FITC | BD Bioscience |
| CD8 | APC-H7 | BD Bioscience |
| CD14 | PE-Cy7 | BD Bioscience |
| CD16 | APC | BD Bioscience |
| CD19 | PerCP | BD Bioscience |
| CD20 | APC-H7 | BD Bioscience |
| CD24 | FITC | BD Bioscience |
| CD25 | BV421 | BD Bioscience |
| CD27 | FITC | BD Bioscience |
| CD27 | BV421 | BD Bioscience |
| CD28 | APC | BD Bioscience |
| CD38 | APC | BD Bioscience |
| CD45 | AF700 | Biolegend |
| CD45 | APC-H7 | BD Bioscience |
| CD45R0 | PE-Cy7 | BD Bioscience |
| CD56 | PE | BD Bioscience |
| CD57 | BV421 | BD Bioscience |
| CD69 | FITC | BD Bioscience |
| CD127 | AF647 | BD Bioscience |
| CD197 (CCR7) | PE | BD Bioscience |
| HLA-DR | V500 | BD Bioscience |
| IgD | PE-Cy7 | BD Bioscience |

## Notes

### Competing Interest Statement

The authors have declared no competing interest.

## References

[1] F.P. Polack, S.J. Thomas, N. Kitchin, et al, “Safety and Efficacy of the BNT162b2 mRNA Covid-19 Vaccine” New England Journal of Medicine, 2020, pp.2603–2615.

[2] M. Voysey, L.Y. Weckx, A.M. Collins, et al, “Single-dose administration and the influence of the timing of the booster dose on immunogenicity and efficacy of ChAdOx1 nCoV-19 (AZD1222) vaccine: a pooled analysis of four randomised trials” The Lancet (British edition), 2021, pp.881–891.

[3] U. Sahin, A. Muik, I. Vogler, et al, “BNT162b2 vaccine induces neutralizing antibodies and poly-specific T cells in humans” Nature, 2021, pp.572–577.

[4] J.R. Barrett, S. Belij-Rammerstorfer, C. Dold, et al, “Phase 1/2 trial of SARS-CoV-2 vaccine ChAdOx1 nCoV-19 with a booster dose induces multifunctional antibody responses” Nature medicine, 2021, pp.279–288.

[5] E.E. Walsh, R.W. Frenck, A.R. Falsey, et al, “Safety and Immunogenicity of Two RNA-Based Covid-19 Vaccine Candidates” New England Journal of Medicine, 2020, pp.2439–2450.

[6] C.B. Jackson, M. Farzan, B. Chen and H. Choe, “Mechanisms of SARS-CoV-2 entry into cells” Nature reviews. Molecular cell biology, 2022, pp.3–20.

[7] H. Gruell, K. Vanshylla, P. Tober-Lau, et al, “mRNA booster immunization elicits potent neutralizing serum activity against the SARS-CoV-2 Omicron variant” Nat Med, 2022, pp.477.

[8] W.T. Harvey, A.M. Carabelli, B. Jackson, et al, “SARS-CoV-2 variants, spike mutations and immune escape” Nature reviews. Microbiology, 2021, pp.409–424.

[9] Anonymous “Mapping the antigenic diversification of SARS-CoV-2” Medical Letter on the CDC & FDA, 2022, pp.88.

[10] M. Hoffmann, N. Krüger, S. Schulz, et al, “The Omicron variant is highly resistant against antibody-mediated neutralization: Implications for control of the COVID-19 pandemic” Cell, 2022, pp.447-456.e11.

[11] E.G. Levin, Y. Lustig, C. Cohen, et al, “Waning Immune Humoral Response to BNT162b2 Covid-19 Vaccine over 6 Months” New England Journal of Medicine, 2021, pp.e84.

[12] F. Muecksch, Z. Wang, A. Cho, et al, “Increased Potency and Breadth of SARS-CoV-2 Neutralizing Antibodies After a Third mRNA Vaccine Dose” bioRxiv: the preprint server for biology, 2022.

[13] D. Geers, M.C. Shamier, S. Bogers, et al, “SARS-CoV-2 variants of concern partially escape humoral but not T cell responses in COVID-19 convalescent donors and vaccine recipients” Sci. Immunol., 2021.

[14] Y. Gao, C. Cai, A. Grifoni, et al, “Ancestral SARS-CoV-2-specific T cells cross-recognize the Omicron variant” Nat Med, 2022, pp.472.

[15] G. Guerrera, M. Picozza, S. D’Orso, et al, “BNT162b2 vaccination induces durable SARS-CoV-2-specific T cells with a stem cell memory phenotype” Science immunology, 2021, pp.eabl5344.

[16] J. Quandt, A. Muik, N. Salisch, et al, “Omicron BA.1 breakthrough infection drives cross-variant neutralization and memory B cell formation against conserved epitopes” Sci. Immunol., 2022.

[17] L. Ruhl, I. Pink, J.F. Kühne, et al, “Endothelial dysfunction contributes to severe COVID-19 in combination with dysregulated lymphocyte responses and cytokine networks” Sig Transduct Target Ther, 2021.

[18] J. Barros-Martins, S.I. Hammerschmidt, A. Cossmann, et al, “Immune responses against SARS-CoV-2 variants after heterologous and homologous ChAdOx1 nCoV-19/BNT162b2 vaccination” Nature medicine, 2021, pp.1525–1529.

[19] M. Becker, A. Dulovic, D. Junker, et al, “Immune response to SARS-CoV-2 variants of concern in vaccinated individuals” Nature communications, 2021, pp.3109.

[20] A. Sokal, M. Broketa, G. Barba-Spaeth, et al, “Analysis of mRNA vaccination-elicited RBD-specific memory B cells reveals strong but incomplete immune escape of the SARS-CoV-2 Omicron variant” Immunity, 2022, pp.1096-1104.e4.

[21] A. Ssemaganda, H.M. Nguyen, F. Nuhu, et al, “Expansion of cytotoxic tissue-resident CD8+ T cells and CCR6+CD161+ CD4+ T cells in the nasal mucosa following mRNA COVID-19 vaccination” Nat Commun, 2022.

[22] T. Schmidt, V. Klemis, D. Schub, et al, “Cellular immunity predominates over humoral immunity after homologous and heterologous mRNA and vector-based COVID-19 vaccine regimens in solid organ transplant recipients” American Journal of Transplantation, 2021, pp.3990–4002.

[23] M.J. van Gils, A. Lavell, K. van der Straten, et al, “Antibody responses against SARS-CoV-2 variants induced by four different SARS-CoV-2 vaccines in health care workers in the Netherlands: A prospective cohort study” PLoS medicine, 2022, pp.e1003991.

[24] E. Lafon, M. Jäger, A. Bauer, et al, “Comparative analyses of IgG/IgA neutralizing effects induced by three COVID-19 vaccines against variants of concern” Journal of allergy and clinical immunology, 2022, pp.1242-1252.e12.

[25] R. Markewitz, D. Juhl, D. Pauli, et al, “Kinetics of the Antibody Response to Boostering With Three Different Vaccines Against SARS-CoV-2” Front. Immunol., 2022.

[26] G.M.N. Behrens, J. Barros-Martins, A. Cossmann, et al, “BNT162b2-boosted immune responses six months after heterologous or homologous ChAdOx1nCoV-19/BNT162b2 vaccination against COVID-19” Nat Commun, 2022.

[27] J. Lang-Meli, H. Luxenburger, K. Wild, et al, “SARS-CoV-2-specific T-cell epitope repertoire in convalescent and mRNA-vaccinated individuals” Nat Microbiol, 2022, pp.675.

[28] A. Tarke, C.H. Coelho, Z. Zhang, et al, “SARS-CoV-2 vaccination induces immunological T cell memory able to cross-recognize variants from Alpha to Omicron” Cell, 2022, pp.847-859.e11.

[29] A. Cossarizza, H. Chang, S. Abrignani, et al, “Guidelines for the use of flow cytometry and cell sorting in immunological studies (third edition)” European journal of immunology, 2021, pp.2708–3145.

